# Bioprocess Modeling Made Easy With Hybrax

**DOI:** 10.64898/2026.11.05.764695

**Authors:** Julian Libiseller-Egger, Mathias Gotsmy, Gonzalo Guillén-Gosálbez, Jürgen Zanghellini

## Abstract

Hybrid models that embed neural networks in ordinary differential equations are increasingly popular in bioprocess modeling, but they are challenging to set up. Much of the effort goes into the code around the model: wet-lab data must be imported and pre-processed, feeds and sampling must be accounted for in the balance equations, and a training loop with cross-validation must be set up. Researchers usually write this code from scratch for each project. In addition, no widely adopted format for bioprocess data exists, and datasets are mostly published in bespoke spreadsheets or not at all. As a result, models and datasets from the literature are difficult to reproduce, reuse, or compare. We present `Hybrax`, an open-source Python framework that addresses these problems. It combines a flexible format for bioprocess data with a customizable end-to-end training pipeline for mechanistic and hybrid models. `Hybrax` derives the transport terms in the process dynamics automatically and provides the utilities needed for training, so that researchers can focus on developing the model itself. It also facilitates model sharing and reproducibility: each training checkpoint contains the model parameters, code, configuration, and training data required to run the model or retrain it from scratch. We demonstrate `Hybrax` by re-implementing two models from the literature, one trained on real-world *E. coli* data and one on synthetic HEK293 data, with performance comparable to the original models. `Hybrax` is available at https://github.com/hybrax/hybrax.

## 1 Introduction

Biotechnological processes are the basis of production for a wide variety of compounds across the pharma-ceutical, chemical, and food industries. Due to high experimental costs and the large design space of possible process operating conditions, the development, optimization, and control of these processes increasingly rely on *in silico* models (Kroll et al. 2017; Helleckes et al. 2023). However, accurately describing bioprocess dynamics remains challenging. Although variables such as temperature, pH, dissolved oxygen, and off-gas composition can be monitored continuously, the underlying physiological state of the production organism is only partially observable (Udugama et al. 2020; Brunner et al. 2021). Hybrid Models (HMs) have been shown to perform well in this data-rich but information-poor domain (Mahanty 2023; Herrera-Ruiz et al. 2025). They are usually implemented as a system of Ordinary Differential Equations (ODEs) in which one or more parameters are estimated by an Artificial Neural Network (ANN), effectively combining the interpretability and physical constraints of a mechanistic model with the flexibility of purely data-driven approaches.

HMs yield robust results, but their setup is not trivial. Model development requires code for importing cultivation data where measurements may have been collected on different time grids, process inputs can be continuous or event-based, and changes in reactor volume due to feeding, sampling, pH control, and potentially evaporation need to be taken into account. Furthermore, successful HM training depends on careful choices of scaling, regularization, early stopping, cross-validation, and other hyperparameters.

Existing open-source tools cover only parts of this workflow. pyFOOMB, for example, supports object-oriented representation, simulation, and parameter estimation of ODE models (Hemmerich et al. 2021), but lacks machinery for efficient ANN training. Other tools offer better integration of ANNs embedded in ODEs: PEtab SciML (Persson et al. 2026) allows solver-independent specification of hybrid models and is supported by PEtab.jl (Persson et al. 2025) and AMICI (Fröhlich et al. 2021). SiNDAE (Lueg et al. 2026), jaxkineticmodel (van Lent et al. 2025), and Catalax (Range 2026) provide related functionality. However, these tools target systems biology, biocatalysis, or scientific machine learning more broadly. Bioprocess-specific features such as feeding, sampling, or multi-stage processes need to be encoded manually for every dataset, if they are supported at all. HYBpy (Pedreira et al. 2025), a Python successor to the MATLAB toolbox HYBMOD (Pinto et al. 2022), does cover bioprocess applications like fed-batch cultivations and offers a user-friendly web application for HM training, but its ease-of-use comes at the cost of flexibility: only a subset of model architectures and no custom loss functions, for instance, are supported. Further, commercial tools such as the Hybrid Modeling Toolbox (Novasign GmbH) and DataHowLab (Data-How AG) contain bioprocess-oriented HM workflows, but are proprietary and closed-source.

Currently, no open-source framework handles the routine parts of setting up, training, and evaluating bioprocess HMs while staying flexible enough for new model designs. Instead, researchers repeatedly spend considerable effort on data cleaning as well as rewriting training code, rather than developing the model itself. Models are typically trained with custom, one-off pipelines (e.g., Ramos et al. 2024; Gotsmy and Guillén-Gosálbez 2026; Martínez et al. 2026) and most data, if published at all, are published in bespoke spreadsheet formats. Applying a model from the literature to one’s own data or vice versa frequently requires a substantial amount of work, impeding study reproducibility and performance comparisons. These issues compound when the number of tested models increases, making comprehensive benchmarks – comparing a meaningful portion of published model architectures on a diverse set of publicly available data – exceedingly difficult to conduct.

Here, we present `Hybrax`, a chassis for bioprocess hybrid modeling that combines a versatile, ontology-inspired representation of bioprocess data with a flexible end-to-end training pipeline for mechanistic and hybrid models. `hybrax.format` is an expressive data structure for bioprocesses of all common types, encoding process configuration (inflows, outflows, media composition, etc.), measurement data, and (optionally) a mathematical process model. From the process configuration, it derives the dilution and feed terms of the process dynamics automatically. Once a dataset has been parsed into `hybrax.format`, HMs can be set up and trained with little additional code. `hybrax.train` provides the training machinery, including data pre-processing, augmentation, and handling of discontinuities such as bolus feeds, allowing researchers to focus on developing the actual process model itself. It employs a modern Universal Differential Equation (UDE) (Rackauckas et al. 2021) approach based on differentiable ODE solvers (Chen et al. 2018), enabling training on concentration data directly without the need for intermediate rate estimation.

In addition to facilitating model development, `Hybrax` also streamlines model sharing. Every training checkpoint contains, alongside the model parameters, all relevant information required to immediately run the model or to fully re-train it from scratch, including pre-processed training data, configuration files, and custom code for the model definition. `Hybrax` is available at https://github.com/hybrax/hybrax.

## 2 Implementation

`Hybrax` is implemented in Python and relies on JAX (Bradbury et al. 2018), Equinox (Kidger and Garcia 2021), and Diffrax (Kidger 2021). Agentic coding tools were used to generate or debug parts of the codebase. Detailed documentation can be found at https://hybrax.readthedocs.io.

The `Hybrax` workflow consists of multiple stages (Fig. 1) forming two conceptual subunits. First, experimental measurements are imported into our novel bioprocess data format using `hybrax.format` and are validated for completeness. Then, the HM is trained via the `hybrax.train` pipeline. Users define the data-driven component (e.g., an ANN) in `custom.py` Python modules. The command-line entry points `prepare` and `train` handle additional data preprocessing (e.g., data augmentation) and training of a single model, while `loo` performs cross-validation and `forward` can be used to evaluate an existing model on a set of initial states and controls.

**Fig. 1.**
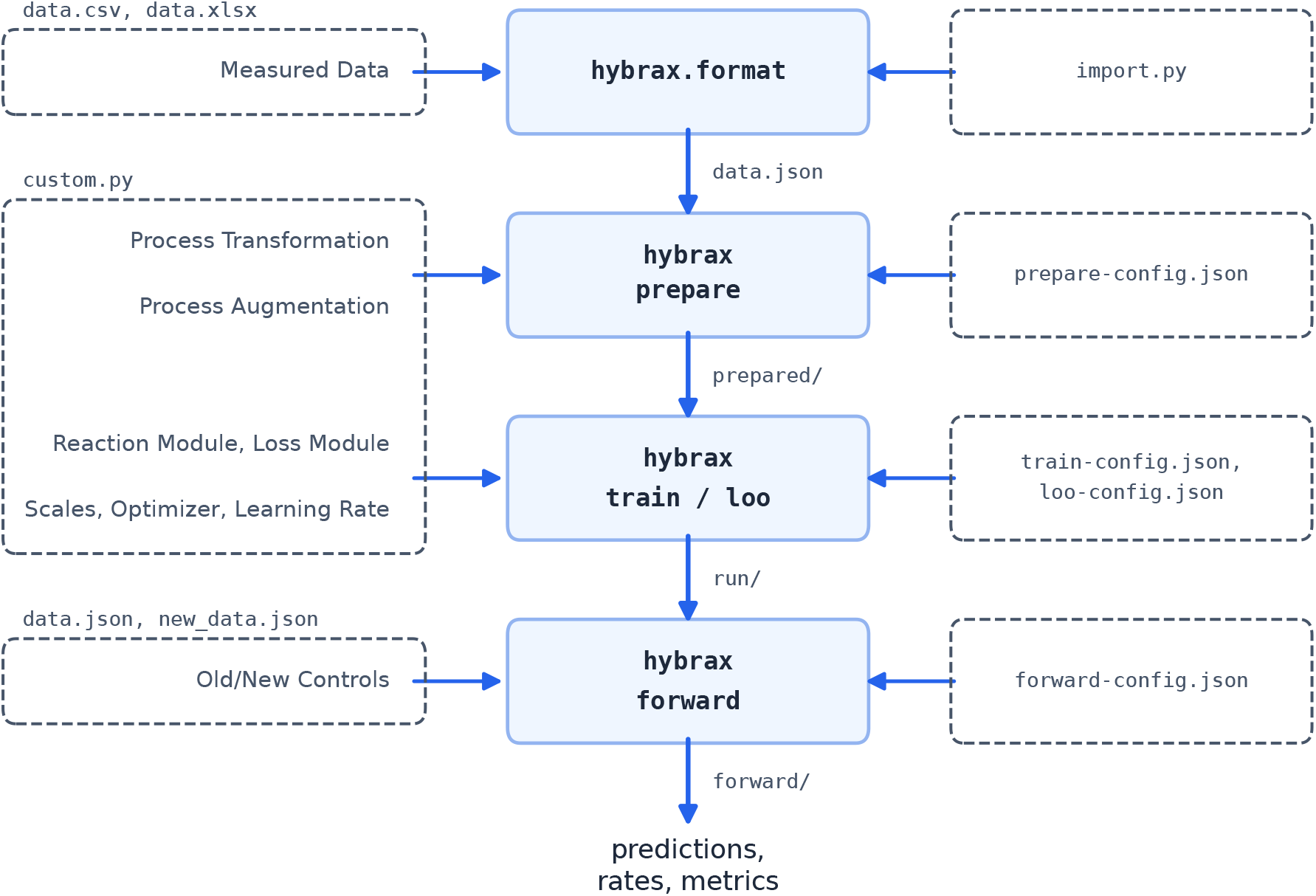
The `Hybrax` user experience: raw measurements are imported once into `hybrax.format`, then move through `hybrax prepare`, `train` (or `loo`), and `forward`, which all can be customized with config files and user-supplied Python code.

All models in `Hybrax` share the same basic structure. The rate of change of the states **y**(*t*) (concentrations and modeled process variables) given the controls **u**(*t*) (feed rates or temperature, etc.) is the sum of a reaction term **R** and a transport term **T**:

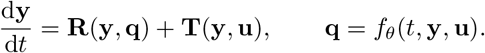

The transport term describes how feeding, sampling, and other volume changes (continuous or discontinuous) alter the concentrations. It is fully determined by the process setup, so `hybrax.format` creates it automatically. The reaction term describes all other process dynamics (usually biological conversions). It is defined by the user (or falls back to a default) in terms of rates **q**, which are computed by *f*_*θ*_. This function can be a set of kinetic expressions, a machine learning model, or a mix of both. `hybrax.train` then fits the parameters *θ* to the data.

### 2.1 hybrax.format

A `hybrax.format` dataset is a `BioProcessCollection` of one or more `BioProcess` objects, typed dataclass trees covering metadata, a shared time axis, reactor volume, reactor-medium composition, process variables, discrete events, and an optional user-defined `ReactionOde`. Every measured or controlled quantity is classified by its physical role: `ReactorMediumComponent` objects represent concentration states (e.g., of biomass, substrate, product) governed by the ODE and by `VolumeChange` objects. A `ProcessVariable` (e.g., pH, temperature) can be a controlled input or a modeled ODE state, but it receives no automatic transport contributions from volume changes.

Volume has its own category in the data model and evolves during ODE integration according to the specified inflows and outflows. Each inflow carries a fixed, user-defined feed-medium composition, while outflow composition follows the reactor medium. When volume and feed compositions are known for a process dataset, `hybrax.format` automatically constructs the transport contribution to the ODE, relieving users of the need to re-derive the corresponding mass balance terms by hand for every model. `Inflow` objects (continuous or bolus feed, base addition for pH control, etc.) increase the reactor volume, whereas `Outflow` objects (sampling, harvest, etc.) reduce it. Continuous outflows carry a `retention` parameter per `ReactorMediumComponent` which can be used to represent evaporation (`retention=1.0` for all components) or perfusion (e.g., `retention=0.99` for biomass, `0` for other components).

Reaction dynamics are generated automatically for each `ReactorMediumComponent`, assuming biomassspecific rates, and for any modeled (i.e., not controlled) `ProcessVariable`. Users can override the default dynamics by supplying symbolic expressions in the `ReactionOde`. For example, if a product accumulates inside the cells, the measured biomass concentration *X* also includes the product *P* . Only the remaining active biomass *X*_*a*_ = *X − P* grows and forms product. In the code, *X*_*a*_ is declared as an algebraic quantity and the reaction terms become *R*_*X*_ = (*q*_*X*_ + *q*_*P*_ ) *X*_*a*_ and *R*_*P*_ = *q*_*P*_ *X*_*a*_, where the rates *q*_*X*_ and *q*_*P*_ are supplied by *f*_*θ*_ later. The transport terms of the ODE are added automatically. With unbounded rates, this is specified in Python as:

~~~
reaction_ode = hybrax.format.ReactionOde(
    algebraic={“Xa”: “X - P”},
    *# ‘(None, None)’ means unbounded rates*
    rates={“q_X”: (None, None), “q_P”: (None, None)},
    derivatives={“X”: “(q_X + q_P) * Xa”, “P”: “q_P * Xa”},
)
~~~

Overriding the default `ReactionOde` in this way encodes prior knowledge of the process dynamics alongside the measured data in a single `hybrax.format` artifact, but it omits the function that estimates the rates. This split is intentional. The effect of intracellular product, in this example, is a fundamental property of the process. *f*_*θ*_, the model that estimates the rates *q*_*X*_ and *q*_*P*_, is not. The resulting decoupled structure makes model comparisons straightforward. In order to benchmark multiple model architectures on the same dataset, researchers only need to swap out the code defining *f*_*θ*_. No further changes to the training pipeline are required.

Data reuse and model benchmarking across studies are further supported by validation functionality provided for `hybrax.format` datasets. Basic validation checks individual process objects for internal consistency and structural problems (e.g., inconsistent volume-change signs, mistimed measurement/sampling alignment). Optional “pre-publication” validation has additional, stricter checks, covering consistent variable definitions, units, and time-axis conventions across a process collection.

### 2.2 hybrax.train

`hybrax.train` exposes four command-line entry points: `prepare`, `train`, and `loo` correspond to the main phases of the training workflow while `forward` uses a trained model to create predictions. Each of these commands loads user-supplied configuration and code files, allowing for extensive customization of the pipeline.

The `prepare` command allows users to perform any data transformations that are required for training but should not modify the raw dataset, such as filtering, data augmentation, or fitting smoothing splines. It also produces diagnostic plots, records provenance metadata, and validates that the prepared dataset is ready for training. We decided to turn `prepare` into its own command to ensure a clean separation between raw data import and training-specific pre-processing. It packages the train-ready data into a single file that researchers can share easily to compare model architectures on identical inputs.

`train` and `loo` both consume this artifact. `train` fits an individual model to the data, whereas `loo` implements cross-validation to train an ensemble (using leave-one-out by default, but the cross-validation strategy is flexible). Users can define models in `custom.py` files, alongside hooks that are executed by `Hybrax` at multiple points in the training process, exposing the loss function, scaling, learning rate schedule, optimizer configuration, etc. to customization. Models must be implemented as Equinox (Kidger and Garcia 2021) modules that calculate the unknown quantities appearing in the ODE right-hand side (`q_X` and `q_P` in our example above). `train` relies on Diffrax (Kidger 2021), which allows for training on concentration data directly without intermediate rate estimation.

Self-contained checkpoints are written to disk at a regular cadence during training. By default, each checkpoint bundles the prepared dataset, model parameters, optimizer state, run configuration, and `custom.py` files. Therefore, they can easily be used for prediction on new data (using the `forward` command), fine-tuning, or retraining from scratch on the same prepared dataset. They also streamline model sharing and benchmarking.

## 3 Results

The documentation at https://hybrax.readthedocs.io provides a variety of usage examples for `Hybrax`, covering multiple mechanistic and hybrid model architectures trained on real-world and synthetic data. Below, we describe three of those examples: a mechanistic model of a microbial process (inspired by Schlögel et al. 2024), a simple ANN-based HM using the same data, and a Long Short-Term Memory (LSTM) HM of a process using HEK293 cells (inspired by Ramos et al. 2024; synthetic data only).

### 3.1 OptFed in Hybrax

Schlögel et al. (2024) created OptFed, a mechanistic model for an *E. coli* process (originally described in Kittler et al. 2022), by first deriving specific ratesv from splines of the concentration data. They then fitted expressions for substrate uptake, maintenance, and product formation to the rates, starting from a larger model and iteratively pruning parameters that did not significantly contribute to the fit. To illustrate how `Hybrax` can also be used for mechanistic models, we fitted the pruned OptFed model directly to the raw concentration data. While not recovering the same final parameter values, the model fitted with `Hybrax` achieved comparable or better agreement with the concentration measurements. Across twelve processes, pooled *R*^2^ values were 0.86 for biomass and 0.85 for product, compared with 0.83 and 0.82 for the original OptFed fit evaluated at the same measurement points. For the product-to-biomass ratio (the optimization target in their study), *R*^2^ was 0.82 versus 0.73. It should be noted that these values describe training-set agreement rather than cross-validation or test-set performance.

### 3.2 Swap the mechanistic for a hybrid model

Once a dataset has been imported into `hybrax.format`, model structures can be swapped out easily. To illustrate this, we reused the dataset from above to train a simple HM where an ANN predicts substrate uptake and the ratios of substrate that are spent on maintenance, product formation, and growth. Despite its small size (2 *×* 4 dense layers, 59 trainable parameters), the model learned the dynamics in the data sufficiently well so that its reconstruction loss is similar to the OptFed kinetic model (*R*^2^ of 0.90 and 0.85 for biomass and product). Again, it is important to emphasize that this is an in-sample result and serves only as an example for the utility of `Hybrax`, not as a direct comparison against the mechanistic model.

### 3.3 An LSTM HM on HEK293 data

The previous example employed a simple feed-forward neural network, but `Hybrax` also supports more complex architectures. To demonstrate this, we implemented an LSTM-based HM for a HEK293 cells process inspired by the work of Ramos et al. (2024).

One challenge of modeling mammalian cell culture compared to microbial processes is that this usually requires tracking considerably more states (concentrations of extracellular amino acids and other metabolites). Consequently, the data-driven component of a HM for a mammalian process needs to predict more rates, making the model more complex. Ramos *et al*. alleviated this issue by transforming all measured concentrations of their dataset into cumulative reacted amounts and then performing Principal Component Analysis (PCA) on the resulting matrix. This decomposition into the most relevant reaction patterns greatly reduced the number of specific rates to be learned by the ANN in their HM formulation. For the nine synthetic cultures available in the authors’ public repository, four principal components retained more than 99% of the variation, so four rates were sufficient to drive the 25 concentration balances. We implemented this approach in `Hybrax`, using a simple continuous-time LSTM rather than the authors’ custom recurrent-cell equations, but keeping their PCA methodology.

As in the authors’ example, five of the nine cultures were used for training, one for validation and checkpoint selection, and three for testing. The selected checkpoint achieved Normalized Root Mean Squared Error (NRMSE) of 0.0567 on validation and 0.0348 on testing, similar to 0.0576 and 0.0376 for the authors’ MATLAB model. One limitation of both implementations is that they do not enforce non-negative concentrations and that the predicted traces therefore sometimes dip below zero. However, `Hybrax` makes it easy to add a penalty to the loss function for such bounds violations, which we used to train a second model with similar performance (0.0582 and 0.0375 validation and test NRMSE) and fewer negative con-centrations. On the test cultures, the average amount by which its predictions fell below zero was 0.04% of each state’s maximum, compared with 0.15% without the penalty and 0.13% for the MATLAB model. Fig. 2 shows the test set predictions of three of the 25 states (biomass, antibody, glucose) for the three models.

**Fig. 2.**
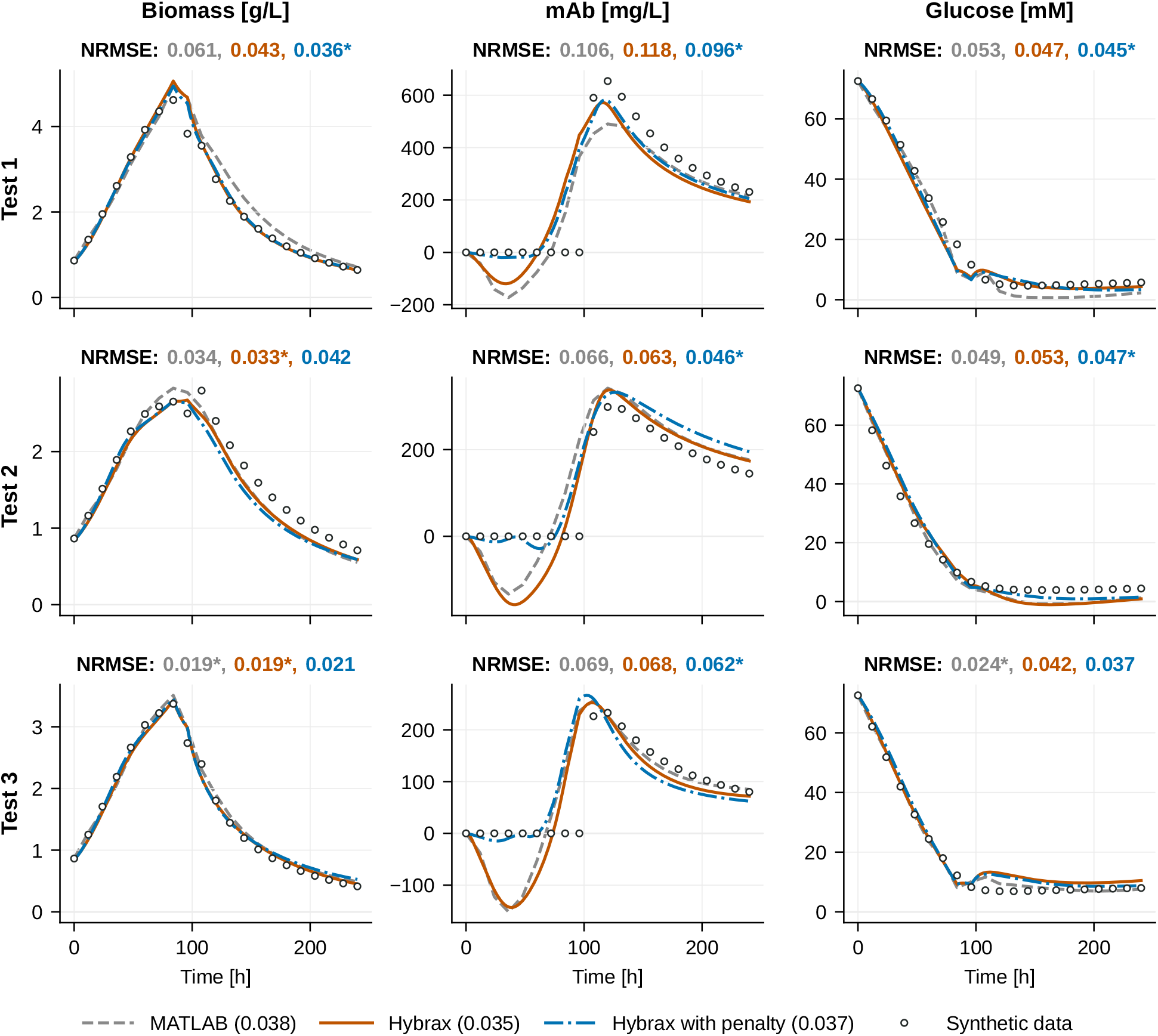
Predictions of biomass, antibody (mAb), and glucose for the three test cultures of the synthetic HEK293 benchmark. The original MATLAB model by Ramos *et al*. is shown in gray and the `Hybrax` models (without and with a penalty on negative concentrations) in orange and blue, respectively. The numbers above each panel give each model’s NRMSE for that culture and state. An asterisk marks the lowest value. The legend gives each model’s NRMSE over all 25 states of the three test cultures. Overall, the three models perform similarly, and the penalty reduces negative concentrations.

## 4 Conclusion

Compared to other areas of computational biology, such as genomics or proteomics, there is considerably less momentum behind the open-source movement in bioprocess modeling. This causes limited interoper-ability of published methods and data, hamstrung benchmarking, and duplicated work, slowing down progress in the field. `Hybrax` represents a unified way of encoding and sharing bioprocess data as well as a feature-rich and customizable pipeline for training ODE-based models with little code.

By handling most of the surrounding functionality, such as ODE transport terms, scaling, and the training loop, `Hybrax` lets researchers focus on the process-specific part of the model. It also provides reusable building blocks and complete examples in the documentation, covering different model types, data augmentation, parameter freezing, and constraint or smoothness penalties. The combination of robust primitives and a diverse set of templates creates a solid foundation for rapid model development and prototyping. In our experience, this is especially relevant in the era of coding agents, which benefit greatly from worked examples.

We will continue to develop `Hybrax` going forward, with a focus on training performance and stability, additional utilities for streamlined data preprocessing, interpretability of prediction results, and uncertainty quantification.

## Statements and Declarations

### Funding

This project was funded in part by the Austrian Science Fund (FWF), grant DOI 10.55776/COE17 – Cluster of Excellence: Circular Bioengineering.

For open access purposes, the authors have applied a CC BY public copyright license to any author-accepted manuscript version arising from this submission.

### Ethics approval and consent to participate

Not applicable.

### Consent for publication

Not applicable.

### Data availability

The *E. coli* data were originally created by Kittler et al. (2022) and published as part of the OptFed modeling study (Schlögel et al. 2024). They are available at https://github.com/gschloegel/OptFed/tree/7811f7b9ef76d624f53b852c03d06cb1b5cc96a3/data.

The data HEK293 created by example Ramos uses et synthetic (2024), al. available at https://github.com/jrcramos/Hybrid-modeling-of-bioreactor-with-LSTM/tree/d4197ed84ba46bfb53c3e4f42cfe00f44916edb5/data.

Both datasets, imported into `hybrax.format`, are also available at https://github.com/hybrax/hybrax.

### Materials availability

Not applicable.

### Code availability

All source code for `Hybrax` as well as the usage examples described in this manuscript are available at https://github.com/hybrax/hybrax under the MIT license. An archived version with a persistent DOI will be deposited in Zenodo upon acceptance.

### Author contributions

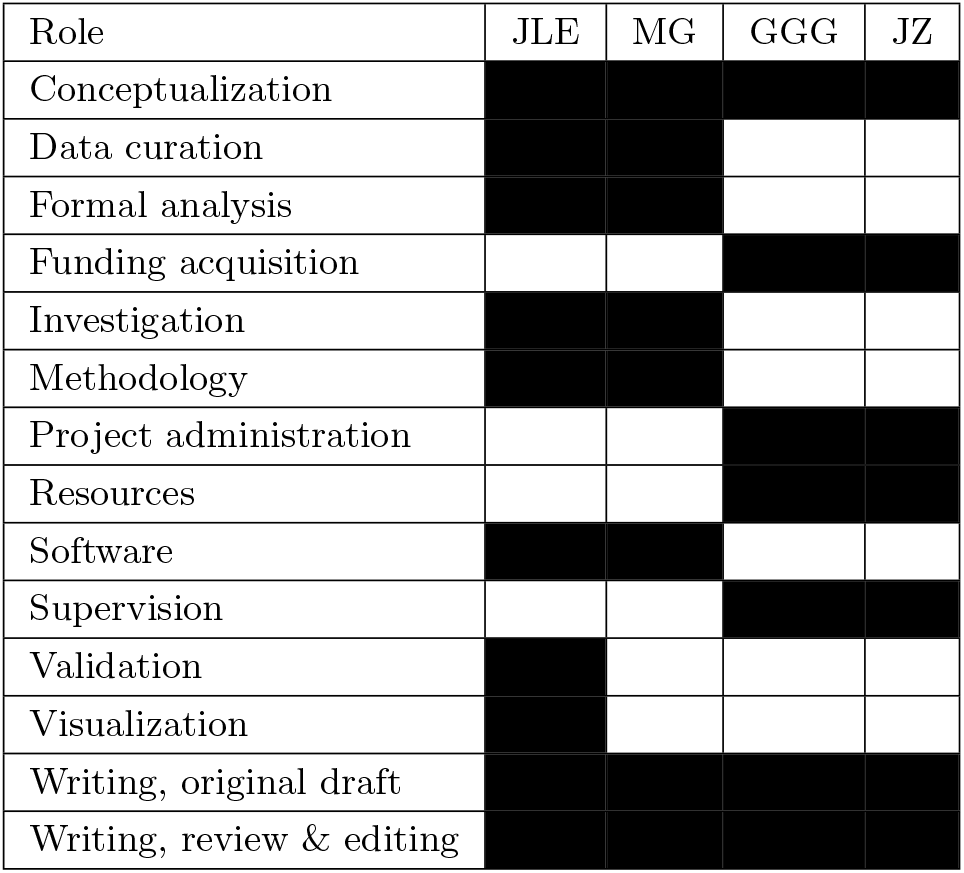

## References

Bradbury, J., Frostig, R., Hawkins, P., Johnson, M.J., Leary, C., Maclaurin, D., Necula, G., Paszke, A., VanderPlas, J., Wanderman-Milne, S., Zhang, Q.: JAX: composable transformations of Python+NumPy programs (2018). https://github.com/jax-ml/jax

Brunner, V., Siegl, M., Geier, D., Becker, T.: Challenges in the Development of Soft Sensors for Bioprocesses: A Critical Review. Frontiers in Bioengineering and Biotechnology 9 (2021) 10.3389/fbioe.2021.722202

Chen, R.T.Q., Rubanova, Y., Bettencourt, J., Duvenaud, D.: Neural Ordinary Differential Equations (2018). https://arxiv.org/abs/1806.07366

Fröhlich, F., Weindl, D., Schälte, Y., Pathirana, D., Paszkowski, L ., Lines, G.T., Stapor, P., Hasenauer, J.: AMICI: high-performance sensitivity analysis for large ordinary differential equation models. Bioinformatics 37(20), 3676–3677 (2021) 10.1093/bioinformatics/btab227

Gotsmy, M., Guillén-Gosálbez, G.: Integrating Metabolic Networks into Hybrid Bioprocess Models. bioRxiv (2026). 10.64898/2026.04.22.720062

Helleckes, L.M., Hemmerich, J., Wiechert, W., von Lieres, E., Grünberger, A.: Machine learning in bioprocess development: from promise to practice. Trends in Biotechnology 41(6), 817–835 (2023) 10.1016/j.tibtech.2022.10.010

Hemmerich, J., Tenhaef, N., Wiechert, W., Noack, S.: pyFOOMB: Python framework for object oriented modeling of bioprocesses. Engineering in Life Sciences 21(3–4), 242–257 (2021) 10.1002/elsc.202000088

Herrera-Ruiz, J.F., Fontalvo, J., Prado-Rubio, O.A.: Hybrid modeling for bioprocesses: Architectures, applications, and perspectives. Engineering Reports 7(12), 70502 (2025) 10.1002/eng2.70502

Kidger, P.: On Neural Differential Equations. PhD thesis, University of Oxford (2021)

Kidger, P., Garcia, C.: Equinox: neural networks in JAX via callable PyTrees and filtered transformations. Differentiable Programming workshop at Neural Information Processing Systems 2021 (2021)

Kittler, S., Ebner, J., Besleaga, M., Larsbrink, J., Darnhofer, B., Birner-Gruenberger, R., Schobesberger, S., Akhgar, C.K., Schwaighofer, A., Lendl, B., et al.: Recombinant protein L: Production, purification and characterization of a universal binding ligand. Journal of Biotechnology 359, 108–115 (2022) 10.1016/j.jbiotec.2022.10.002

Kroll, P., Hofer, A., Ulonska, S., Kager, J., Herwig, C.: Model-Based Methods in the Biopharmaceutical Process Lifecycle. Pharmaceutical Research 34(12), 2596–2613 (2017) 10.1007/s11095-017-2308-y

Lueg, L.R., Alves, V., Schicksnus, D., Kitchin, J.R., Laird, C.D., Biegler, L.T.: A simultaneous approach for training neural differential-algebraic systems of equations. Computational Optimization and Applications (2026) 10.1007/s10589-026-00823-y

Mahanty, B.: Hybrid modeling in bioprocess dynamics: Structural variabilities, implementation strategies, and practical challenges. Biotechnology and Bioengineering 120(8), 2072–2091 (2023) 10.1002/bit.28503

Martínez, C., Rocha Calvette, F., Péré, M., Barrientos, M., Ossandón, S.: Hybrid modeling framework for bioprocesses with minimal prior knowledge and limited data. bioRxiv. Version 2, posted 2026-01-11 (2026). 10.1101/2025.11.14.688550

Pedreira, J., Pinto, J., Gonçalves, D., Barahona, P., Oliveira, R., Costa, R.S.: HYBpy: A web-based framework for hybrid modeling of biological systems. Digital Chemical Engineering 17, 100278 (2025) 10.1016/j.dche.2025.100278

Persson, S., Fröhlich, F., Grein, S., Loman, T., Ognissanti, D., Hasselgren, V., Hasenauer, J., Cvijovic, M.: PEtab.jl: advancing the efficiency and utility of dynamic modelling. Bioinformatics 41(9), 497 (2025) 10.1093/bioinformatics/btaf497

Persson, S., Snelling, B., Philipps, M., Weindl, D., Cvijovic, M., Hasenauer, J., Pathirana, D., Fröhlich, F.: PEtab SciML: an exchange format for specifying and training dynamic scientific machine learning models (2026). https://arxiv.org/abs/2608.20184

Pinto, J., Mestre, M., Ramos, J., Costa, R.S., Striedner, G., Oliveira, R.: A general deep hybrid model for bioreactor systems: Combining first principles with deep neural networks. Computers & Chemical Engineering 165, 107952 (2022) 10.1016/j.compchemeng.2022.107952

Rackauckas, C., Ma, Y., Martensen, J., Warner, C., Zubov, K., Supekar, R., Skinner, D., Ramadhan, A., Edelman, A.: niversal Differential Equations for Scientific Machine Learning (2021). https://arxiv.org/abs/2001.04385

Ramos, J.R.C., Pinto, J., Poiares-Oliveira, G., Peeters, L., Dumas, P., Oliveira, R.: Deep hybrid modeling of a HEK293 process: Combining long short-term memory networks with first principles equations. Biotechnology and Bioengineering 121(5), 1554–1568 (2024) 10.1002/bit.28668

Range, J.: Catalax: a JAX-based framework for (neural) ODE modelling in biocatalysis. GitHub. Version 0.5.5 (2026). https://github.com/JR-1991/Catalax Accessed 2026-10-22

Schlögel, G., Lück, R., Kittler, S., Spadiut, O., Kopp, J., Zanghellini, J., Gotsmy, M.: Optimizing bio-processing efficiency with OptFed: Dynamic nonlinear modeling improves product-to-biomass yield. Computational and Structural Biotechnology Journal 23, 3651–3661 (2024) 10.1016/j.csbj.2024.09.024

Udugama, I.A., Gargalo, C.L., Yamashita, Y., Taube, M.A., Palazoglu, A., Young, B.R., Gernaey, K.V., Kulahci, M., Bayer, C.: The Role of Big Data in Industrial (Bio)chemical Process Operations. Industrial & Engineering Chemistry Research 59(34), 15283–15297 (2020) 10.1021/acs.iecr.0c01872

van Lent, P., Bunkova, O., Magyar, B., Planken, L., Schmitz, J., Abeel, T.: Jaxkineticmodel: Neural ordinary differential equations inspired parameterization of kinetic models. PLOS Computational Biology 21(7), 1012733 (2025) 10.1371/journal.pcbi.1012733

